# Salicaceae endophyte inoculation alters stomatal patterning and improves the intrinsic water-use efficiency of *Populus trichocarpa* after a water-deficit

**DOI:** 10.1101/2024.08.10.607466

**Authors:** Matthew Hendrickson, Darshi Banan, Robert Tournay, Jonathan D. Bakker, Sharon L. Doty, Soo-Hyung Kim

## Abstract

Microorganisms may enhance plant resilience to water stress by influencing their hosts’ physiology and anatomy at the leaf-level. Bacterial and yeast endophytes, isolated from wild poplar and willow, can improve the intrinsic water-use efficiency (*iWUE*) of cultivated poplar (*Populus*) under water-deficits by lowering stomatal conductance (*g_sw_*). However, the relevance of stomatal anatomy underlying this reduction remains unclear. We hypothesized endophyte inoculation could change host stomatal anatomy, and this would relate to decreases in *g_sw_*. We subjected Salicaceae endophyte-inoculated and uninoculated *Populus trichocarpa* to well-watered and water-deficit treatments in greenhouse studies. We examined the changes of individual stomatal traits and related the composition of these parameters, termed stomatal patterning, to leaf gas-exchange under light saturation. After a water-deficit, inoculation improved *iWUE* at light saturation from preserving carbon assimilation (*A_net_*) and lowering *g_sw_*, but these changes were independent of soil-moisture status. Drops in *g_sw_* corresponded to underlying shifts in stomatal patterning. Inoculated plants had smaller, more compact stomata and greater anatomical maximum stomatal conductance (*g_smax_*) relative to the control. Salicaceae endophytes may alter stomatal density and size, lowering *g_sw_* and increasing *iWUE*. Future efforts may quantify endophyte colonization of the host to draw direct relationships between microbes and stomatal traits.

**HIGHLIGHT:** Poplars inoculated with specific bacteria had leaves containing many, tiny pores relative to the trees without the microbes; these plants with the small, dense pores related to greater intrinsic water-use efficiency.

## INTRODUCTION

Climate change is extending regional droughts, threatening the sustainable productivity of vulnerable forest systems (IPCC 2023; Vose et al. 2016). To preserve stand growth without increasing irrigation, silviculture industries can improve the water-use efficiency (*WUE*) of trees (Stenzel et al., 2021). *WUE* represents the ratio of carbon gained for biomass to water lost and can be applied at scales ranging from entire ecosystems down to individual leaves (Hatfield & Dold, 2019). At the leaf-level, *WUE* focuses on extrinsic and intrinsic processes. Extrinsic *WUE* (*WUE*) relates net instantaneous rate of carbon assimilation (*A_net_*) to leaf transpiration (*E*), and thus indirectly accounts for environmental variables through their effects on E. In contrast, intrinsic *WUE* (*iWUE*) relates *A_net_* to stomatal conductance to water vapor (*g_sw_*) and thus depends solely on the inherent physiology of a species (Rho & Kim, 2017).

Commercially and ecologically valuable trees such as black cottonwood (*Populus trichocarpa)* are susceptible to water stress, and increasing *WUE* remains a target to maintain their yields (Marron et al., 2014). While refining the *WUE* of these trees has relied on genomic-assisted breeding, this approach is slow due to *Populus* reproduction rates compared to environmental change (Stanton et al., 2014; Biselli et al., 2022). To supplement this process, induced acclimation can leverage the phenotypic plasticity of plant leaves to modify *iWUE* (Flexas et al., 2013, Leakey et al., 2019). Altering *iWUE* requires adjustments to the stomatal pores embedded in the leaf epidermis, as these link the intercellular space to the environment. Functionally, these openings balance the exchange of internal water vapor loss with aerial carbon assimilation by responding to external and inner signals (Harrison et al., 2019; Buckley 2019). Plants control this gaseous tradeoff by dynamically shifting the aperture of existing pores or altering the size, number, and distribution of pores in newly emergent leaves (Lawson et al., 2014).

Stomatal adjustments can influence other physiological aspects such as evaporative leaf cooling (Urban et al., 2017), mass-flow nutrient uptake (He & Dijkstra, 2014), and biomass accumulation (Afas et al., 2006). Consequently, adaptations of stomatal traits to a prolonged drought evokes a cost-benefit effect, incurring a biological gain at the expense of another (Muir, 2015; de Boer et al., 2016; Mckown et al., 2019). This presents a challenge to boost *iWUE* of cultivated trees like black cottonwood that may inherently tailor stomatal patterning to maximize growth rates instead of water conservation under dry conditions (Dunlap & Stettler, 2001; Pearce et al., 2006; Mckown et al., 2014, 2019). Therefore, refining *iWUE* in poplars may require innovative methods, and one flexible strategy applies the mutualistic partnership between plants and microorganisms (Doty et al., 2017; Rho et al., 2018c).

A plant and its microbiome form a holobiont, or discrete ecological unit, coadapted to a specific environment (Vandenkoornhuyse et al., 2015). In its native riparian habitat along the North American Cascade Range, black cottonwood carries distinct strains of endophytes – microbes that live inside plants - that vary with climatic locations (Firrincieli et al., 2020; Doty et al., 2009). Select bacterial endophytes assist black cottonwood by fixing dinitrogen gas in return for photosynthates or inorganic acids (Doty et al. 2016; Rho et al., 2018a; Kandel et al. 2015). When strains like these are isolated and used to inoculate other plant species, they provide diverse host benefits such as greater biomass, seedling viability, and improved water relations (Khan et al., 2012; Aghai et al., 2019, Rho et al., 2018c; Rho et al. 2020a). While endophytes offer plant growth promotion (PGP) and resilience to water stresses, an effect on a particular PGP mechanism is highly context dependent (Rho et al., 2018b).

Plant-microbe-environment interactions are complex (Rho & Kim, 2017). For example, studies of the effects of Salicaceae-derived endophytes under water-stress report varied effects. A recent study showed that inoculated black cottonwood that were exposed to water-stress had improved *iWUE* because *A_net_* was maintained while midday *g_sw_* decreased (Banan et al., 2024). Water-stress experiments in species such as hybrid poplar (Khan et al., 2016) and rice (Rho et al., 2018c) have revealed comparable results. However, under different stresses, endophytes show no effect on *g_sw_* (Knoth et al., 2014; Rho et al., 2020a, Rho et al., 2020b), and the greater literature details mixed outcomes on *g_sw_* based on diverging mechanisms (Rho & Kim 2017). Variation of this highly dynamic parameter across studies encourages an examination of the morphological traits that underly its physiology. For example, Rho et al. (2018c) attributed their observed reduction in *g_sw_* partly to decreases in abaxial stomatal density.

Plants may identify beneficial endophytes through microbe-associated molecular patterns (MAMPs) as a part of innate immunity (Carvalho et al. 2016). Although the connection between MAMPs and stomatal closure has received extensive study through the concept of stomatal defense, the importance of these signals on stomatal development needs further investigation (Melotto et al., 2017). Microbes can interact with a plant through immune receptors, secondary messengers, and hormones. For instance, Salicaceae endophytes can generate plant hormones (Khan et al., 2016), and have shown to increase abscisic acid (ABA) concentration in leaf tissue (Rho et al., 2018c) – a crucial regulator of stomatal dynamics and development. While a theory exists of pathogenic influence on leaf gas-exchange via shifts in stomatal morphology (Muir, 2020), the empirical relationship of commensal endophytes on these mechanisms remains unclear. Particularly, the extent of PGP endophytes on the anatomical stomatal traits underlying *g_sw_* has received little attention (Rho et al., 2018c; Rho et al. 2020b). This study examined the impact of Salicaceae endophytes and water stress on stomatal morphology in black cottonwood and related stomatal morphology to leaf gas-exchange.

### Research questions and hypotheses

We performed a factorial greenhouse experiment to determine the influence of endophyte inoculation and water stress on the gas exchange and stomatal patterning of black cottonwood. We defined stomatal patterning as the multivariate set of anatomical stomatal traits (**Table 1**). The rationale for the hypotheses was based on the understanding that certain Salicaceae endophytes produce hormones such as ABA that could affect stomatal development (Khan et al., 2016, Rho et al., 2018c). Therefore, the hypotheses were to determine whether inoculation of *Populus trichocarpa* with specific Salicaceae endophytes would: (1) lower operational *g_sw_* while maintaining the maximum assimilation rate (*A_max_*) under saturating light conditions, (2) change stomatal patterning, and (3) coordinate operational *g_sw_* with stomatal patterning. Plants with and without inoculation were exposed to water-deficit or kept well-watered conditions to determine if inoculation conferred an advantage during water stress.

**Table 1.** List of parameters referenced within text and the accompanying units.

| Parameter | Definition | Units |
| --- | --- | --- |
| GCL | Guard cell length. Distance of the stomatal complex on the major axis from the outermost edges of the guard cells. Measured for both adaxial (Ad) and abaxial (Ab) leaf sides. | $\mu\text{m}$ |
| $g_{\text{smax}}$ | Maximum anatomical stomatal conductance. Latent variable determined from geometric assumptions that describes the greatest conductance based on stomatal anatomy. Calculated for both adaxial (Ad) and abaxial (Ab) leaf sides. | $\text{mol m}^{-2} \text{s}^{-1}$ |
| PL | Pore length. Distance of the stomatal pore on the major axis from the innermost edges of the guard cells. Measured for both adaxial (Ad) and abaxial (Ab) leaf sides. | $\mu\text{m}$ |
| PW | Pore width. Distance of the stomatal pore on the minor axis from the innermost edges of the guard cells. Measured for both adaxial (Ad) and abaxial (Ab) leaf sides. | $\mu\text{m}$ |
| SD | Stomatal density. Number of stomata per unit area. Measured for both adaxial (Ad) and abaxial (Ab) leaf sides. | $\text{mm}^{-2}$ |
| SI | Stomatal index. Number of stomata per total number of epidermal cells (pavement cells + stomata). Measured for both adaxial (Ad) and abaxial (Ab) leaf sides. | - |
| SR | Stomatal ratio. Number of adaxial (Ad) stomata per abaxial (Ab) stomata. | - |
| $A_{\text{max}}$ | Light saturated $A_{\text{net}}$ under $2000 \mu\text{mol}_{\text{photon}} \text{m}^{-2} \text{s}^{-1}$ of photosynthetic photon flux density (PPFD) at ambient $[\text{CO}_2]$ at $25^\circ\text{C}$ | $\mu\text{mol m}^{-2} \text{s}^{-1}$ |
| $A_{\text{net}}$ | Net $\text{CO}_2$ assimilation rate. | $\mu\text{mol m}^{-2} \text{s}^{-1}$ |
| $E$ | Transpiration, or water lost through stomata in response to environmental conditions | $\text{mmol m}^{-2} \text{s}^{-1}$ |
| $g_{\text{sw}}$ | Stomatal conductance to water vapor. | $\text{mol m}^{-2} \text{s}^{-1}$ |
| $i\text{WUE}$ | Intrinsic water-use efficiency. Leaf-level trait that describes the ratio of $A_{\text{net}}$ to $g_{\text{sw}}$ | $\mu\text{mol mol}^{-1}$ |
| $\text{WUE}$ | Extrinsic water-use efficiency. Leaf-level trait that describes the ratio of $A_{\text{net}}$ to $E$ | $\mu\text{mol mol}^{-1}$ |

Each stomatal trait contains a prefix of Ad or Ab to correspond it to either the adaxial (upper) or abaxial (lower) leaf surface, respectively. Shaded cells represent variables that encompass stomatal patterning (i.e., composition of stomatal traits); unshaded cells indicate leaf gas-exchange parameters.

## MATERIALS AND METHODS

### Endophyte origin and inoculum preparation

The consortium consisted of nine endophyte strains, eight bacteria and one fungus, isolated from *Populus* and *Salix* species (**Table 2**; termed DOE mix). Strains were selected based on their PGP traits and ability to improve host plant growth under nitrogen-limitation (Knoth et al., 2014), drought (Khan et al., 2016), or for their synergistic microbe-microbe interactions (Sher et al., in review). The annotated draft genomes for the newly published strains have been deposited in the NCBI GenBank database under BioProject PRJNA1058978, while the strains for *R. aceris* WP5, *Rhizobium* sp. PTD1, and *Rhodoturula graminis* WP1 were previously reported under the BioProjects PRJNA247595, PRJNA247570, and PRJNA32711, respectively.

**Table 2.** List of endophyte strains used in the consortium.

| Endophyte | Strain | Host source | Biosample | Reference |
| --- | --- | --- | --- | --- |
| *Azotobacter beijerinckii* | SD1 | *P. trichocarpa* | SAMN40215788 | This study |
| *Azospirillum* sp. | 11R-A | *P. trichocarpa* | SAMN39201418 | Sher et al. (in review) |
| *Herbiconiux* sp. | 11R-BC | *P. trichocarpa* | SAMN39201421 | Sher et al. (in review) |
| *Sphingobium* sp. | 11R-BB | *P. trichocarpa* | SAMN39201427 | Sher et al. (in review) |
| *Rahnella aceris* | R10 | *P. trichocarpa* | SAMN40215789 | This study |
| *Sphingomonas* sp. | WW5 | *Salix sitchensis* | SAMN39212602 | Knoth et al. 2014 |
| *Rahnella aceris* | WP5 | *P. trichocarpa* | SAMN02787083 | Doty et al. 2009 |
| *Rhizobium* sp. | PTD1 | *P. trichocarpa x deltoides* | SAMN02797158 | Doty et al. 2005 |
| *Rhodotorula graminis* | WP1 | *P. trichocarpa* | SAMN00138171 | Xin et al. 2009 |

Endophytes were isolated from trees on the Westside of the Cascade Range.

Pure-culture isolates from -80 C cryostocks were streaked on either Mannitol-Glutamate/Luria Bertani (MGL, Cangelosi et al. 1991) or Nitrogen-limited Combined Carbon Media (NL-CCM, Rennie, 1981) agar and incubated at 30 C for 24-48 hr. Isolated colonies were used to inoculate 250 mL of either MGL or NL-CCM broth, and placed on an orbital shaker (150 RPM, 30 C, 24-48 hr.). The cultures were then pelleted via centrifugation (6000 RPM, 10 min), washed with Nitrogen Free Medium (Sher, et al. 2024), and resuspended via vortexing; this process was repeated three times. The cells were suspended in 20-50 mL NFM, the OD600 recorded, then mixed with sterilized 50% glycerol and stored at -80 C until used. The final inoculum was prepared by combining the thawed cryostocks to a final OD600 = 0.28.

Internally-sterile Populus trichocarpa clone Nisqually-1 plants were propagated in Woody Plant Medium (Phytotech) containing 1 µg/ml indole butyric acid rooting hormone. Rooted apical cuttings of similar age were transferred to sterile 1L vessels containing approximately 300 ml sterile DQ water (Millipore) one day prior to co-cultivation to remove adhering agar. Rinsed plants were transferred to 1L vessels containing 100 ml of the washed consortium, and co-cultivated for 2-3 days . Inoculated plants were then transferred to individual sterile beakers containing NFM within sterile Magenta vessels.

### Background and design

The experiment took place in a controlled greenhouse at the Center of Urban Horticulture, University of Washington, and ran from 05/31/2022 to 08/20/2022. A total of 36 *Populus trichocarpa* Nisqually-1 clones were used and half were inoculated with the DOE endophyte consortium 83 days prior to the experiment (**Table 2**). Plants were transplanted in separate 8.5-liter pots with an average of 1940 g of sterile rooting media (5-parts peat: 1-part coarse pumice by weight). After transplanting, the pots were placed under a mist tent to promote rooting for one week before moving them to the greenhouse bench where they acclimated for 50 days before the experimental water-deficit treatment was imposed. Pots were arranged in a randomized complete block design (RCBD) to isolate plant heterogeneity when applying the abiotic treatments. Blocks were perpendicular to the length of the 5.5 m bench to account for potential environmental gradients across the greenhouse. There were 9 blocks in total with one replicate from each treatment group within a block. During the experiment, the plants experienced a 14/10-hour light/dark photoperiod from supplemental LED lamps (ZELION HL300 Grow White, OSRAM Sylvania Inc., Wilmington, MA, USA). Environmental conditions during the experiment can be found in **Table 3**. Plants were fertigated with a 200 ppm 17-5-17 Nutriculture liquid nutrient solution (Plant Marvel Laboratories Inc., Chicago Heights, IL, USA) every 2-3 days. Each plant was placed in a plastic bucket on the greenhouse bench to prevent cross-contamination of endophyte treatments through fertigation run-off Calibrating the experimental water-deficit involved creating a simple linear model estimating soil-moisture level from mass. To create this relationship, we weighed ten additional 8.5-liter pots with the same type and mass of experimental media under water-saturated (100% water holding capacity (WHC)) and oven-dried (0% WHC) conditions. Saturation occurred by evenly wetting the media and covering the tops to minimize water loss from evaporation. All potted media reached a stable mass after two days, and their values were recorded as 100% WHC. Drying ensued by moving the same pots into a forced air oven at 70 °C. After weighing biweekly for 12 days, all pots reached a consistent mass, and their values were logged as 0% WHC.

**Table 3.** Summary table of the greenhouse experiment presented in this paper, highlighting the key environmental data.

| Experiment summary |  |
| --- | --- |
| Water-deficit length | Summer<br>(06/15/2022 - 08/17/2022); 64 days |
| Inoculation date | 03/09/2022 |
| Endophyte consortia | DOE |
| Experimental design | Randomized complete block design; 2x2 factorial |
| Fertigation regime | 17-5-17 NPK at 200 ppm; supplied every 2-3 days depending on pot WHC |
| Average daily light integral* | 47.1 mol m<sup>-2</sup> day<sup>-1</sup> (± 14.4) |
| Average relative humidity | 69.6 % (± 8.75) |
| Average temperature | 21.9 °C (± 1.3) |

Average values of environmental data given (± SD). \*Average daily light integral (DLI; mol<sub>photon</sub> m<sup>-2</sup> day<sup>-1</sup>) represents daily solar irradiance values (MJ m<sup>-2</sup> day<sup>-1</sup>) obtained from a weather station located outside the greenhouse (AgWeatherNet, WSU, 2023). Note the true DLI values plants experienced are different from these values because of the transmission through glazing materials, obstruction by greenhouse structure, and supplemental lighting.

The water-stress treatment was applied by reducing the fertigation for 9 randomly selected inoculated plants and 9 randomly selected control plants. This treatment was imposed 98 days after inoculation and 50 days after the plants were placed on the bench. The soil-moisture deficit was defined from the calibration curve as 35% of the pot WHC; the remaining 18 plants received a well-watered treatment (80% of WHC). Plants in the water-deficit group received fertigation at an increased frequency, attempting to account for potential differences in nutrient availability. Pots were weighed every 2-3 days, and their mass was converted to WHC using the calibration curve. Plant mass was assumed negligible in this calculation. When the mass of a pot fell below its target threshold, it was fertigated back to an upper limit weight (45% WHC for WD; 100% WHC for WW), such that the WHC could oscillate around the target value (**Fig. 1**).

**Figure 1.**
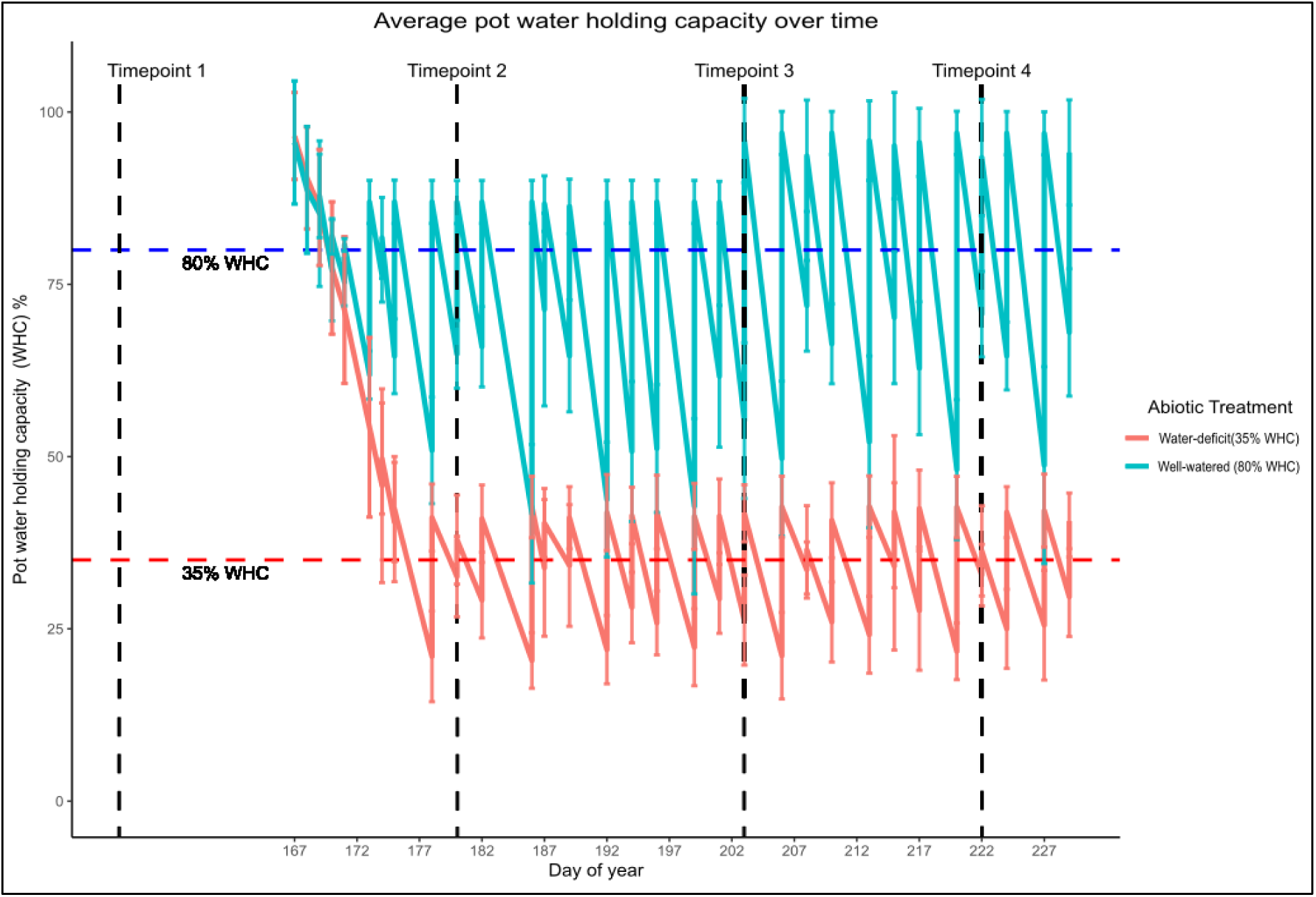
Well-watered (80% WHC) and water-deficit (35% WHC) treatment progression expressed as a percentage of pot water holding capacity (WHC) over time. The blue line indicates the well-watered group, and the red line shows the water-deficit group. Individual pots were weighed every two-three days and rewatered with nutrient solution to their target WHC based on from an initial calibration curve. Vertical dashed lines represent the sampling timepoints when leaf gas-exchange and leaf epidermal impressions were taken.

### Leaf gas-exchange

Leaf gas-exchange measurements were performed with an open-flow, portable photosynthesis system (LI-COR 6400XT; LI-COR, Lincoln, NE, USA), equipped with a 2 cm^2^ chamber head. Sampling occurred on all plants in randomly selected blocks at four sampling timepoints. One sampling took place before the experimental water stress treatment was imposed (t1, n = 4). The remaining sampling occurred while the water-deficit treatment was underway (t2, n = 6; t3, n = 8; t4 n = 9; **Fig. 1**). For each sampled plant, the chamber was clamped to the youngest, fully-expanded, sun-exposed leaf (i.e., 4 – 5 leaves distal to the apical meristem) in the middle of the lamina away from the midvein. Each leaf was wrapped in tinfoil for at least 20 minutes before sampling so that it was dark-adapted initially. Within the cuvette, blue and red light was provided with the 6 cm^2^ LED source (LI-6400-02B) to generate uniform intensity across the examined leaf area. Using a built-in photosynthetic light response program of the LI6400 (*AQ* curves), the photosynthetic photon flux density (PFD) was increased stepwise from 0 μmol_photon_ m^-2^ s^-1^ to 2000. μmol_photon_ m^-2^ s^-1^ while keeping the reference CO_2_ (*C_a_*) at 400 μmol m^−2^. Net CO_2_ assimilation (*A*_net_) was allowed to stabilize to the new chamber condition based on criteria from the auto program at 0, 20, 50, 100, 200, 400, 600, 800, 1000, 1500, 2000 μmol_photon_ m^-2^ s^-1^ (i.e., for 15 seconds, ΔCO2S < 1; ΔH2OS <1; Flow < 1). Constant chamber settings represented ambient environmental conditions on a warm day (leaf temperature, 25°C; relative humidity, 40-60%; flow rate, 500 μmol s^−1^). After completing the preset program, a spot measurement was logged. Diurnal influences were minimized by completing measurements between 8:00-16:00 on the same day.

### Stomatal morphology

Leaf impressions were made with dental putty (Colténe/Whaledent AG, Alstätten, Switzerland) from the top (adaxial) and bottom (abaxial) sides of each plant’s youngest, fully expanded, sun-exposed leaf. Sampling timepoints matched with leaf gas-exchange (**Fig. 1**). For plants in which gas-exchange was measured, putty was dispensed on the same area where these measurements were made. For other plants, putty was dispensed evenly over the middle of the lamina and away from the midvein. In total, 252 leaf impressions were taken over the course of the experiment.

Clear nail varnish was applied on the silicone leaf impressions, allowed to dry, and then removed with clear adhesive tape. After mounting each resulting positive impression onto a microscope slide, images were taken through a microscope (E200 Eclipse, Nikon Instruments Inc., Melville, NY, USA) with a camera attachment (MU-1803 HS, AMScope, Irvine, CA, USA) using a constant image resolution (1782×1372 pixels). Image dimensions were converted to distances by using a calibration ratio created by measuring a stage micrometer with the same image resolution in ImageJ (Schneider et al., 2012). At least three images per impression at every time point were taken under 400x magnification (i.e., 0.068 mm^2^/image) to measure individual stoma. Four images per impression were taken at every time point under 100x magnification (i.e., 1.104 mm^2^/image) to determine stomatal density.

From the images taken at 400x magnification, guard cell length (GCL; µm) and pore length (PL; µm) of 10 stomata were measured across the images for each replicate using ImageJ (Schneider et al., 2012). Pore width (PW) was measured with StomataGSmax (Gibbs et al., 2021) after training and validating a new model based on 122 annotated photos (Epochs = 150; IOU = 0.8; Loss = 0.03; **Fig. S1**). Stomatal index (SI) was measured manually by counting cells with the Cell Counter plugin in ImageJ. Each 100x image was first processed by StomataCounter (Fetter et al., 2019) before manual annotation to determine stomatal density (SD; mm^-2^).

Measurements were averaged across images for each plant and timepoint to avoid pseudoreplication. Anatomical maximum stomatal conductance (*g_smax_*; mol m^-2^ s^-1^) was calculated using the size and density data with the equations of Franks & Farquhar (2001) and geometric assumptions based on Franks & Beerling (2009):

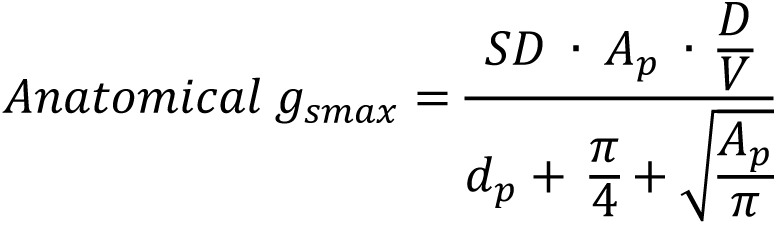

where SD is the average stomatal density, D is the diffusivity of water in air (2.82 × 10−5 m^2^ s^−1^ at 25°C) and *V* is the molar volume of air (m^3^ mol^−1^, at 25°C and 101.3 kPa). Pore depth (*d_p_*; m) was equal to average guard cell width, represented as a quarter of the guard cell length (*d_p_* = GCL/4). The mean maximum stomatal pore area (*A_p_*, m^2^) was calculated assuming stomatal pores were circular at maximum opening (Franks & Beerling, 2009), with the diameter equal to pore length (*A_p_* = (PL/2)^2^). **Fig. 2** displays a schematic representation of how stomatal morphology encompasses patterning.

**Figure 2.**
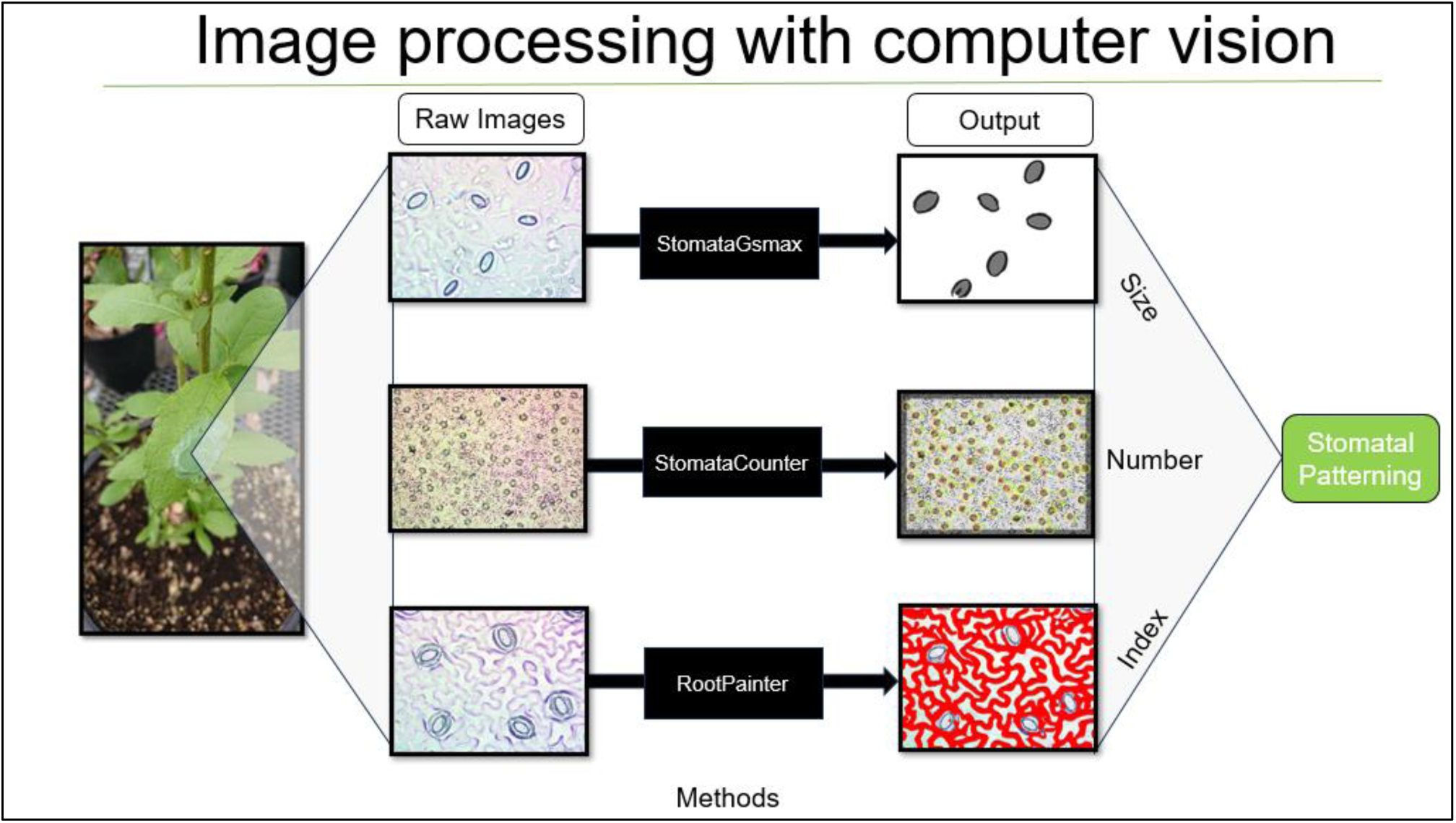
Schematic representation showing the image processing to gather stomatal traits. Images of impressions were taken at 400x magnification for stomatal dimension and index data, and 100x for density. Convolutional neural network (CNN) models were trained and validated from StomataGSmax (Gibbs et al., 2021) for automating morphological measurements and RootPainter (Smith et al. 2022) and StomataCounter (Fetter et al., 2019), to assist manual counting of cells. Any manual measurements were performed in ImageJ.

### Data analysis

#### Leaf gas-exchange

To determine treatment differences for each spot measurement, non-parametric permutational ANOVA tests were used to avoid making assumptions about data normality, equal variance, or sphericity (Frossard & Renaud, 2021). For the pre-water-stress observation (t1), mixed effect models were used to compare each trait between inoculated and control plants. Blocking terms were modeled as random effects to reduce unexplained variability. Post-water-deficit observations also used mixed effects models to compare fixed effects of endophyte treatments, water-stress, timepoint, and two-way interaction terms. In addition to blocks, plants were considered as the random variable to capture the within sample correlation from repeated measures. Post-hoc contrast analyses were examined using Wilcoxon-Mann-Whitney tests for pairwise comparisons. Statistical significance was determined if values were lower than the type I error rate (α = 0.05). Corrections for multiple tests were not made for a priori tests but were made for a posteriori tests via the Bonferroni-Holm corrections. Adjusted Partial eta squared 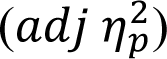 effect sizes were generated for each result to avoid possible overestimations of the effect on the population as follows (Mordkoff 2019):

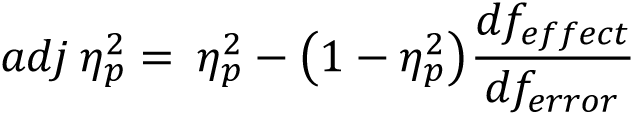

#### Principal component analysis of stomatal morphology

Traits of stomata morphology were highly correlated (**Fig. S9**) and thus were combined through principal component analysis (PCA) into uncorrelated principal components (PCs). Separate PCAs were conducted for pre water-deficit treatment (t1), post water-deficit treatments (t2, t3, t4), and across the entire experiment (t1, t2, t3, t4). Interpretations of PCs were based on the magnitude and sign of the vector associated with each stomatal trait. Though other PCs were also examined, the first two were the most important, and we decided the first two PCs sufficiently captured the patterns in stomatal patterning. Biplots were generated and the inoculation and water-deficit treatments overlaid onto them. Each PC was analyzed using statistical models with the structures described above for gas-exchange spot measurements. This was used as an alternative to performing several hypothesis tests to compare each individual stomatal trait among treatments. However, individual hypothesis tests were performed and analyzed (**Fig. 4**; **Fig S12-13**).

#### Stomatal patterning

Permutational multivariate analysis of variance (PERMANOVA) was used to test treatment effects on stomatal patterning (Anderson 2001; **Table 1**). This was used to avoid making several statistical assumptions and assumed the accumulated difference from individual stomatal traits could reveal a collective difference (Anderson 2017). This differs from the PCA as it encompassed all the variation in stomatal patterning, including that which was not represented in the first two PCs. Prior to the statistical tests, each stomatal trait was rescaled as a proportion of its range (i.e., min-max normalized). Effectively, this preserved the observed relationships within each stomatal trait but gave equal importance across traits. The similarity of stomatal patterning between each plant was determined by generating a Euclidean distance matrix. All post-water deficit observations were aggregated to examine the average treatment effects on stomatal patterning as time and inoculation did not interactively affect individual stomatal traits. Sequential sum of squares (i.e., type I) was used to allow treatment factors placed first in the model explain most of the variation.

#### Corresponding stomatal patterning with conductance

Simple linear regression was used to correspond stomatal patterning with conductance in part to avoid multicollinearity from using individual traits of stomatal anatomy as dependent variables. All post water stress observations were aggregated to simplify the analysis under the justification that the correlation structure of stomata patterning was well preserved over time (**Fig. S9**). We began with a global model which included the two PCs, inoculation and water-stress treatments, and blocking terms. Operational *g_sw_* under light saturation (PFD = 2000 μmol_photon_ m^-2^ s^-1^) was the response. Global models were then reduced by removing the most complex terms, often the interactions. The final model was selected to balance complexity with parsimony and consisted of each PC and the blocking term. Mixed effects models were again used, considering the blocking term as a random effect and the PCs as fixed. Conditional R^2^ was chosen to evaluate the model, which allowed both the PCs and blocking effects to explain the response in stomatal conductance. Maximum likelihood was used to estimate the parameters, which allowed for significance testing of models.

#### Statistical and graphing software

All analyses were performed in R v4.2.2 (R Core Team, 2023), using *tidyverse* functions (Wickham et al., 2019). PERMANOVA was run with the *adonis2* function from *vegan v2.6-4*; univariate parametric and permutational ANOVAs were performed with the *aovperm* function from *permuco v1.1.2*; PCA biplots were created using *ggbiplot v0.55*; correlograms were generated with the *ggcorrplot v0.1.4* package; model selection and exploration was done with *MuMIn v1.47.1*, *lmerTest v3.1-3*, and *lme4 v1.1-33*; confidence intervals and bootstraps were generated with *boot v1.3-28*; nonlinear least squares fitting was performed using *nls.multstart v1.2.0*; violin plots and response curves were created using *cowplot v1.1.1* and *ggh4x v0.2.4*. Inkscape *v1.2.0*, a vector-based imaging editor, was used to increase text size after exporting figures from R.

## RESULTS

### Leaf gas-exchange

Spot measurements of leaf gas-exchange were examined for all treatments. Prior to water stress (t1), endophyte inoculation from the DOE mix significantly reduced the net carbon assimilation rate (*A_net_*) and intrinsic water-use efficiency (*iWUE*) under light saturation (2000 μmol_photons_ m^-2^ s^-1^; (**Table 4**; **Fig. 3**). However, inoculation showed no effect on stomatal conductance (*g_sw_*; p = 0.44). Inoculation marginally impacted leaf transpiration (*E*) and significantly reduced extrinsic water-use efficiency (*WUE*; **Table 4**; **Fig. S4**).

**Table 4.** Effect sizes (adj η^2^) of each treatment for the examined traits before and after water-stress.

|  | *PC1* | *PC2* | *A<sub>net</sub>* | *g<sub>sw</sub>* | *iWUE* | *E* | *WUE* | *T<sub>leaf</sub>* |
| --- | --- | --- | --- | --- | --- | --- | --- | --- |
|  | *Stomatal density and size tradeoff* | *Stomatal ratio* |  |  |  |  |  |  |
|  | Pre-water deficit |  |  |  |  |  |  |  |
| Inoculation | **0.28** | 0.01 | **0.34** | 0 | **0.19** | 0.02* | **0.16** | 0.01 |
|  | Post-water deficit |  |  |  |  |  |  |  |
| Inoculation | **0.10** | 0 | 0 | **0.05** | **0.05** | 0.03* | **0.05** | 0.00 |
| Water-deficit | **0.12** | 0 | **0.24** | **0.25** | **0.12** | **0.24** | **0.08** | **0.06** |
| Inoculation x Water-deficit | 0.01 | 0 | 0 | 0 | 0 | 0 | 0 | 0 |
| Timepoint | 0 | 0 | **0.21** | **0.20** | **0.06** | **0.28** | **0.11** | **0.09** |
| Inoculation x Time | 0.001 | 0 | 0 | 0 | 0.02 | 0 | 0.02 | 0 |
| Water-deficit x Time | 0.02 | 0.02 | **0.07** | **0.08** | 0.01 | **0.09** | 0 | 0 |

Significant terms ( $\alpha = 0.05$ ) are bolded and based on non-parametric ANOVA models. Examined traits include PC1, interpreted as the tradeoff between stomatal density and size and PC2, interpreted as the stomatal density ratio driven by shifts in adaxial numbers. Gas exchange parameters from response curves at $2000 \mu\text{mol}_{\text{photon}} \text{m}^{-2} \text{s}^{-1}$ include net assimilation ( $A_{\text{net}}$ ; $\mu\text{mol m}^{-2} \text{s}^{-1}$ ); operational stomatal conductance ( $g_{\text{sw}}$ ; $\text{mol m}^{-2} \text{s}^{-1}$ ); intrinsic water-use efficiency ( $iWUE$ ; $\mu\text{mol mol}^{-1}$ ); transpiration ( $E$ ; $\text{mol m}^{-2} \text{s}^{-1}$ ); extrinsic water-use efficiency ( $WUE$ ; $\mu\text{mol mol}^{-1}$ ); leaf temperature ( $T_{\text{leaf}}$ , °C). Values labeled with “\*” had marginal significance ( $p < 0.1$ )

**Figure 3.**
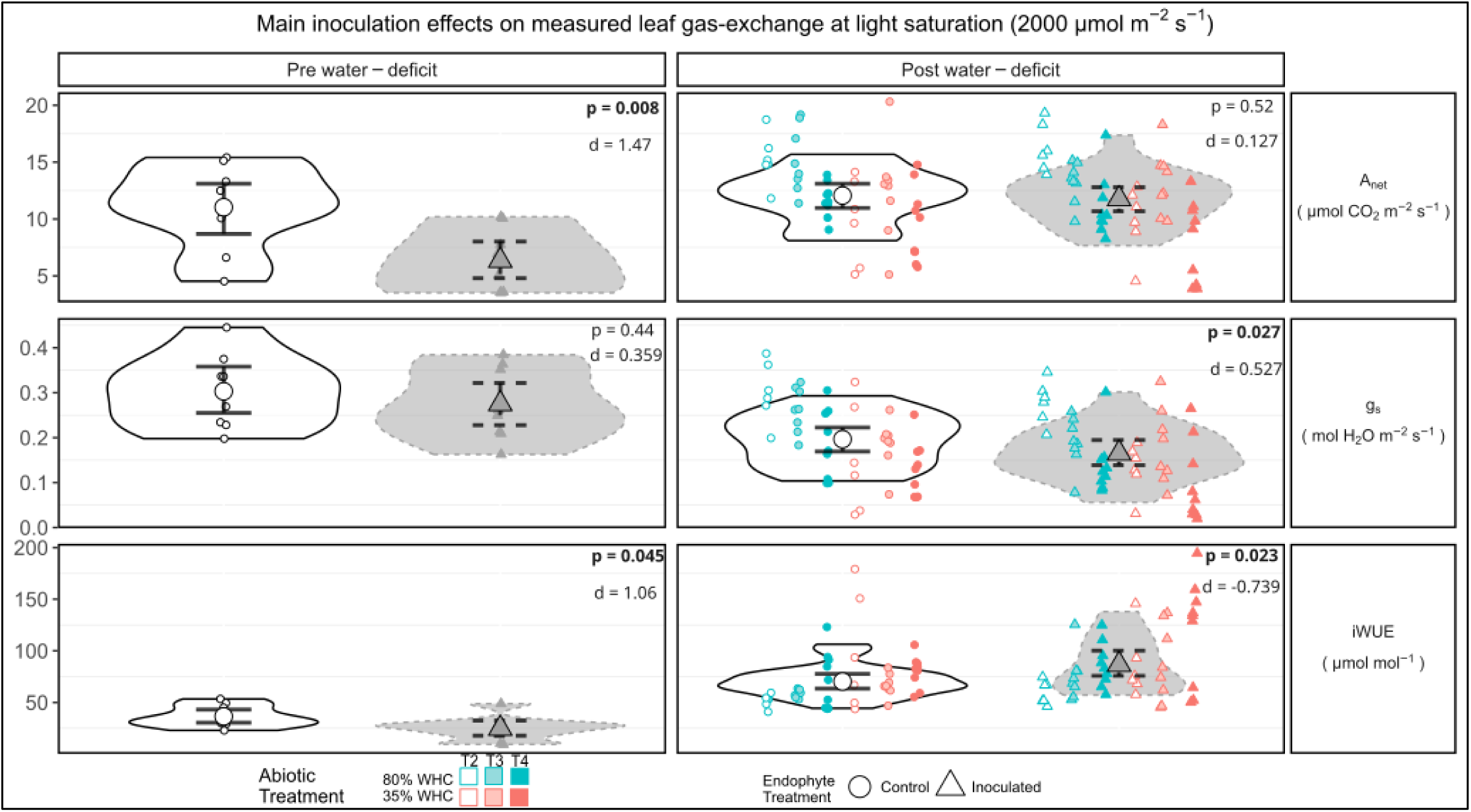
Main inoculation effect on leaf gas-exchange taken at light saturation (2000 μmol_photons_ m^-2^ s^-1^) from the youngest, fully expanded leaf of uninoculated (control; circles) and endophyte-inoculated (triangles) *Populus trichocarpa* pre and post water-deficit. This includes plants from both 80% and 35% WHC groups (blue & red, respectively) at each sampling timepoint (T2, T3, T4). Enlarged shapes represent the mean and lines illustrate the 95% confidence intervals of the net assimilation rate (*A_net_*), stomatal conductance (*g_sw_*), and intrinsic water-use efficiency (*iWUE*). Cohen’s d effect sizes shown for comparisons of inoculation treatments to express the magnitude of mean differences. Confidence intervals were created from the quantiles of case-resampled bootstrap replicates (n = 1000).

After the water-deficit treatment was imposed, inoculation did not affect *A_net_* but lowered *g_sw_* and thus increased *iWUE* (**Table 4**; **Fig. 3**). Inoculation also marginally influenced *E* (p <0.077) and therefore *WUE* (p = 0.04). However, inoculation effects did not interact with time or water-deficit (**Fig. S2**). The water-deficit treatment increased *iWUE*, and over time, it decreased *A_ne_*_t_ and *g_sw_* (**Table 4**; **Fig. 3)**. Likewise, the water deficit increased WUE (p = 0.001) from decreases of *E* over time (p <0.001). Time influenced all leaf gas-exchange parameters (**Table 4**; **Fig. S2**).

### Principal component analysis of stomatal morphology

Most stomatal traits were highly correlated, and these were consistent over time (**Fig. 4**; **Fig. S9**). Therefore, these individual traits could be blended into composite principal components to reveal changes of stomatal patterning. Generally, stomatal size and density traits were inversely correlated, and these traits were orthogonal to stomatal ratio and index. Another observation was that the strength of the relationship between anatomical *g_smax_* and stomatal size diminished post water-deficit.

**Figure 4.**
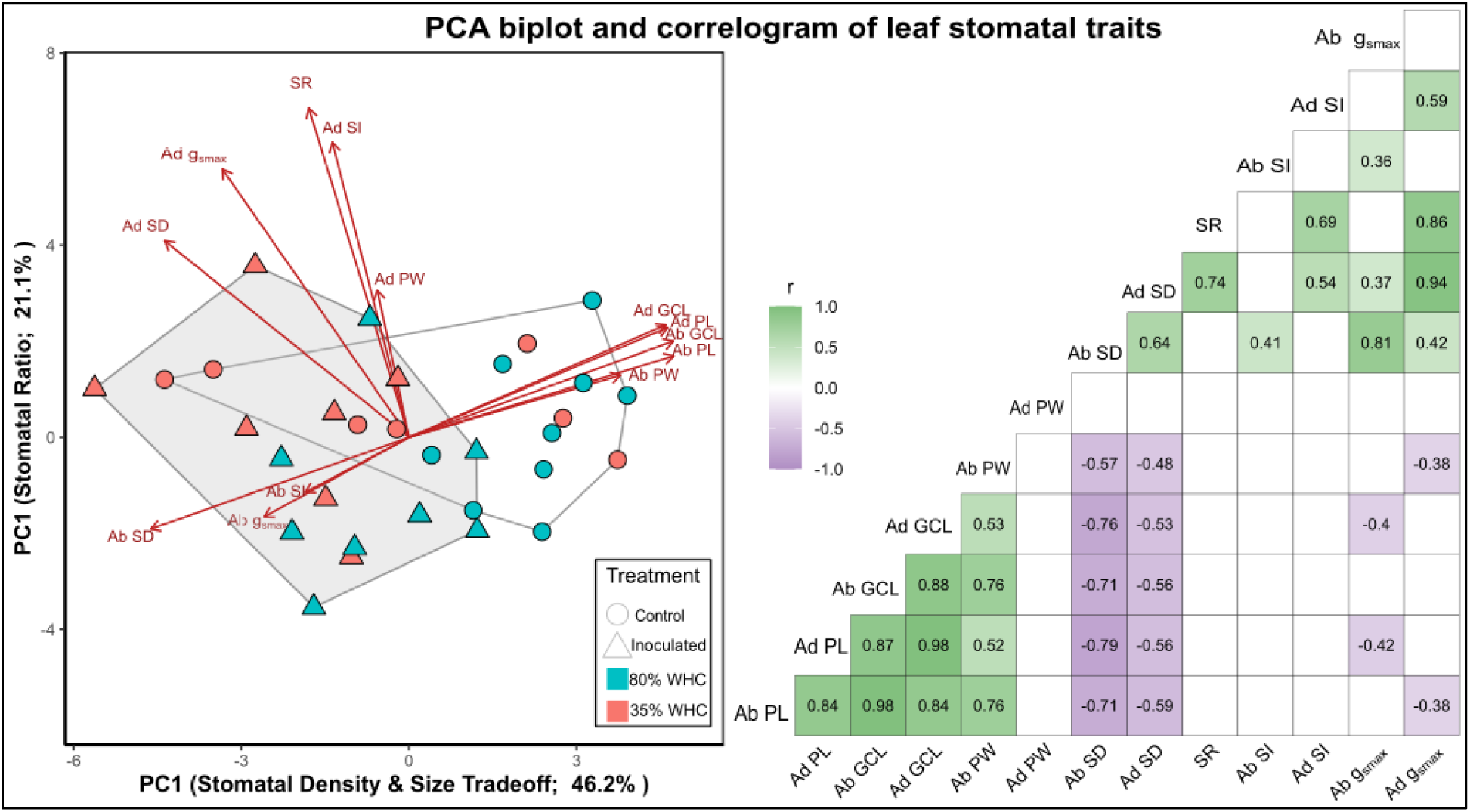
Two-dimensional principal component analysis (PCA) biplot (left) and correlation matrix (right) of stomatal patterning traits in *Populus trichocarpa* aggregated across the experiment. Based on magnitude and direction of vectors, PC1 represents the tradeoff between stomatal density and size; PC2 shows the distribution of stomata to leaf sides. Convex hulls indicate the main endophyte effect with white and grey representing control and inoculated plants, respectively. Pearson’s correlation coefficient (r) is shown for each significant relationship (p <0.05) along with either green or purple colors indicating positive or negative correspondence, respectively. Blank cells represent nonsignificant correlations. For both adaxial (Ad) and abaxial (Ab) leaf surfaces, examined traits include guard cell length (GCL), pore length (PL), pore width (PW), stomatal index (SI), anatomical maximum stomatal conductance (*g*_smax_), stomatal density (SD), and stomatal density ratio (SR).

At each timepoint, the first two principal components (PCs) accounted for more than 58% of the variation (**Fig. S11**). Although the loadings changed at each timepoint, the interpretation of the PCs shared the same understanding; therefore, the PCs were aggregated across the entire experiment (**Fig. S10**). PC1 gave similar weights to all stomatal size traits and bottom stomatal density. Traits that represented stomatal dimensions had opposite signs of those associated with numbers. This was interpreted as the tradeoff between stomatal size and density and accounted for 46.2% of the variance in the stomatal variables. PC2 gave most weight to stomatal ratio and adaxial density, index, and anatomical *g_sma_*_x_, and was interpreted as the distribution of stomata to leaf side, driven by shifts in adaxial stomatal numbers. This accounted for 21.1% of the variation. Along PC1, increasing values represented plants with larger, more sparse stomata, whereas decreasing values depicted plants with denser, smaller stomata. For PC2, increasing values portrayed plants with higher stomatal ratios due to more adaxial stomata (**Fig. 4**).

Prior to water stress (t1), endophyte inoculation from the DOE mix significantly altered the stomatal density and size relationship (PC1; **Table 4**; **Fig. 4**). Inoculated plants tended to have smaller, denser stomata with greater anatomical maximum stomatal conductance (*g_smax_*) relative to uninoculated plants. No effects were seen on the distribution of stomatal numbers to leaf side (PC2). After starting the water-deficit treatment, inoculation continued to affect the stomatal density and size relationship (PC1) in the same way (**Table 4**). Water-deficit also significantly impacted this relationship: plants experiencing water-stress had smaller and denser stomata than those that were not water stressed (**Table 4**; **Fig. 4**). Time did not affect either PC, nor were any interactions significant (**Table 4**). For individual traits, inoculation strongly reduced guard cell and pore lengths on both leaf sides (**Fig. 5**; **S12**).

**Figure 5.**
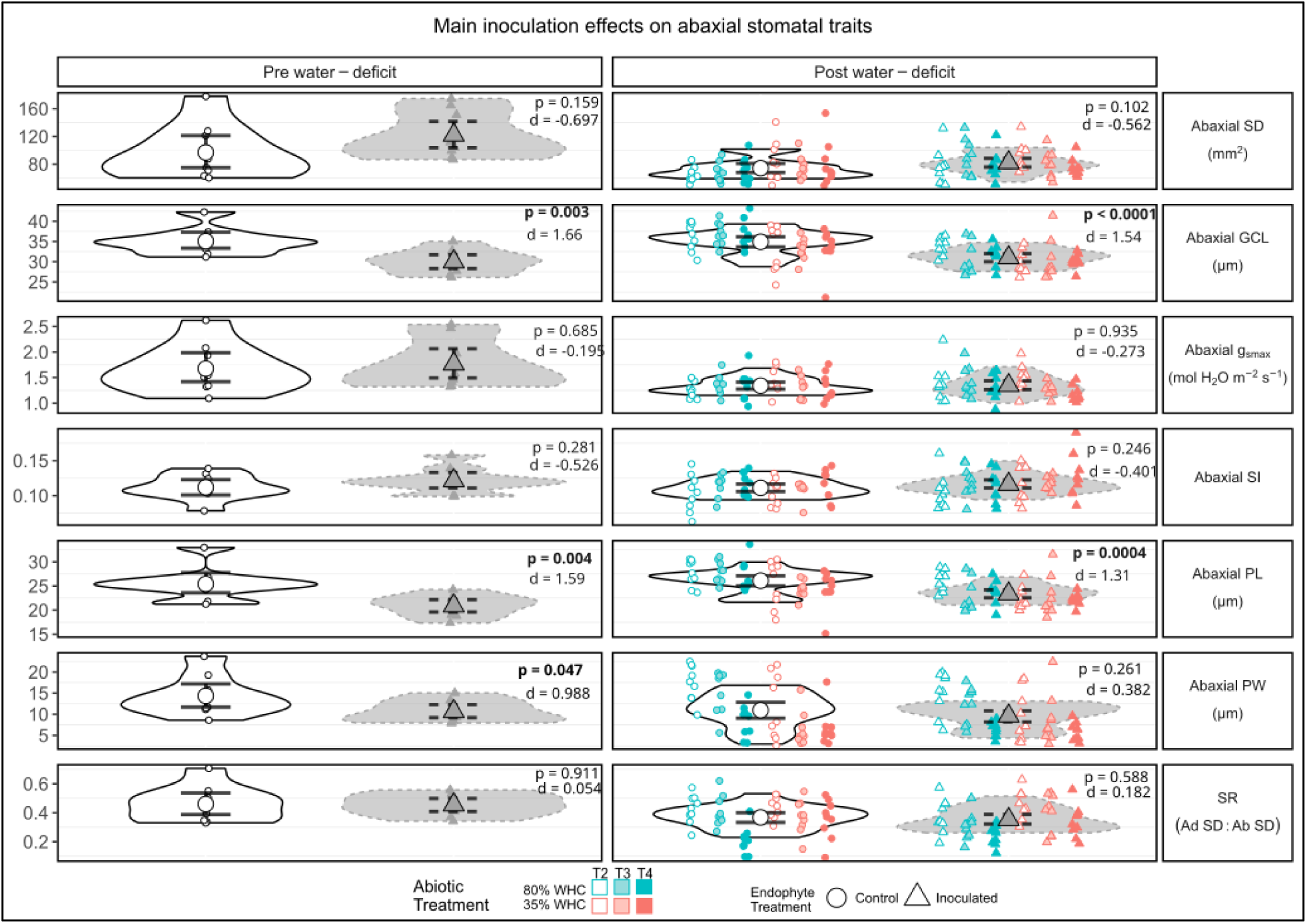
Main inoculation effect on abaxial stomatal traits from the youngest, fully expanded leaf of uninoculated (control; circles) and endophyte-inoculated (triangles) *Populus trichocarpa* pre and post water-deficit. This includes plants from both 80% and 35% WHC groups (blue & red, respectively) at each sampling timepoint (T2, T3, T4). Enlarged shapes represent the mean and lines illustrate the 95% confidence intervals of the guard cell length (GCL), anatomical maximum stomatal conductance (g_smax_), stomatal index (SI), pore length (PL), and pore width (PW). Cohen’s d effect sizes shown for comparisons of inoculation treatments to express the magnitude of mean differences. Confidence intervals were created from the quantiles of case-resampled bootstrap replicates (n = 1000).

### Relating stomatal patterning to conductance

Simple linear regression was used to determine the relationship between the principal components of stomatal traits with operational *g_sw_* under light saturation (PFD = 2000 μmol_photons_ m^-2^ s^-1^). Before water stress, both PC1 (stomatal density & size tradeoff) and PC2 (stomatal ratio, driven by increases in adaxial density) marginally related to *g_sw_* (*R*^2^_*conditional*_ = 0.24.; p = 0.1). After the water-deficit, the model containing PC1 and PC2 corresponded to *g_sw_* and had notable explanatory power when allowed to vary with soil moisture status (*R*^2^_*conditional*_ = 0.63; p = 0.002). While PC1 was a significant predictor within each water level (p <0.001 for both), PC2 only related to *g_sw_* under well-watered conditions (p = 0.04). The fitted relationship between *g_sw_* and PC1 showed a positive relationship, meaning as operational *g_sw_* increased so did the magnitude of the PC scores representing larger, sparser stomata with lower anatomical g_smax_ (**Fig. 5**). PC2 also showed this direct relationship, meaning as operational *g_sw_* increased so did the magnitude of the PC scores representing higher ratios due to increases in adaxial stomatal numbers (**Fig. 6**).

**Figure 6.**
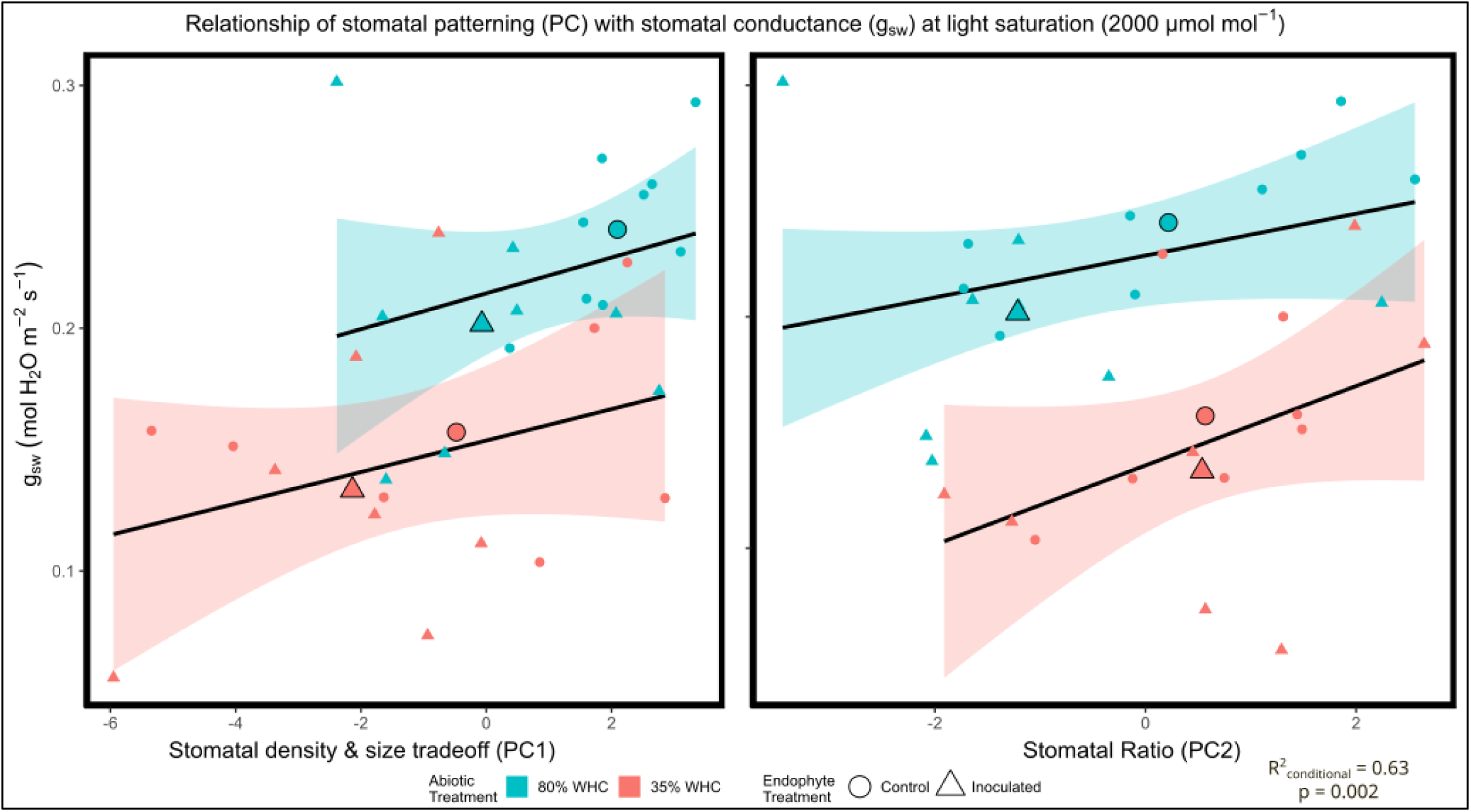
Principal component regression (PCR) illustrating the relationship between stomatal patterning and conductance of uninoculated (circles) and inoculated (triangles) plants under well-watered (80% WHC; blue) and water-deficit (35% WHC; red) conditions. Principal components (PCs) encompass 13 traits relating to stomatal morphology, and operational conductance (*g_sw_*) was taken at light saturation (2000 μmol_photons_ m^-2^ s^-1^). PC1 represents the tradeoff between stomatal density and size – decreasing values indicate smaller and denser stomata with greater anatomical g_smax_; increasing values represent larger, sparser stomata with lower anatomical g_smax_. PC2 depicts shifts in stomatal ratio, driven by changes in adaxial numbers – decreasing values portray lower ratios due to decreases in adaxial stomata, increasing values represent higher ratios from increases in adaxial stomata. Enlarged shapes illustrate the mean value for stomatal patterning and conductance of each treatment (i.e., the group centroid) to assist with interpretation of the trends.

## DISCUSSION

This study explored new questions and mechanisms regarding the influence of specific Salicaceae endophytes on intrinsic water-use efficiency (*iWUE*) in black cottonwood (*Populus trichocarpa*) under a water-deficit that corroborate and expand on previous findings (Banan et al., 2024). Our study showed that the DOE mix enhanced *iWUE* in black cottonwood under high light after imposing a water-deficit. These improvements corresponded to shifts in stomatal patterning, characterized by smaller guard cells and pores. In this study, the notable relationships between compositional stomatal traits and operational *g_sw_* highlighted the importance of morphology, but the unexplained variation implies temporal stomatal dynamics plays an influential role. These empirical findings can guide future efforts which may quantify the microbiome and explore the underlying mechanisms of endophyte-host signaling.

### Endophytes improved iWUE of Populus trichocarpa under high light after a water-deficit

We hypothesized inoculation to improve *iWUE* (*A_net_/g_sw_*) under high light after experiencing a soil-moisture deficit. Our leaf gas-exchange data supported this, showing improvements in *iWUE* at light saturation (**Table 4**; **Fig. 2**), consistent with previous studies that used Salicaceae endophytes in black cottonwood and other species (Banan et al., 2024; Khan et al., 2016; Rho et al., 2018). After the water-deficit, these enhancements were driven by reductions in *g_sw_* and maintenance of *A_net_*. Our results differed than the previous study with black cottonwood as the decreases of *g_sw_* from inoculation did not depend on soil-moisture status after starting the water-deficit (Banan et al., 2024). This physiological response likely depends on specific signals generated by the endophytes of microbes and their interactions with other abiotic factors (Melotto et al., 2017).

One pathway to interpret these observations involves phytohormones. Certain Salicaceae endophyte strains used in this experiment, such as WP1, PTD1, WP5, and WW5 (**Table 2**), have shown to generate abscisic acid (ABA), brassinosteroids (BR), gibberellic acid (GA), indole-3-acetic acid (IAA), jasmonates (JA), and salicylic acid (SA; Khan et al. 2016). These hormones are involved in the complex crosstalk that affects stomatal movement, and apart from IAA and a specific jasmonate (JA-Ile), all generally can promote stomatal closure (Acharya & Assmann, 2009; Melotto et al. 2017). Though black cottonwood and related hybrids have shown limited stomatal sensitivity to ABA (Marron et al., 2014; Ridolfi & Dreyer, 1997), this hormone promotes solute leakage from guard cells (Schulte & Hinckley, 1987). If the endophytes increase leaf ABA content *in vivo*, as seen in rice (Rho et al. 2018c), the higher accumulation may lead to osmotic adjustments of the guard cells over time, reducing stomatal aperture and conductance. Future work may confirm the presence and timing of ABA concentrations within inoculated black cottonwood leaves to further understand the influence of endophyte signaling on phytohormones.

We also observed that inoculated plants experienced a lower *iWUE* at light saturation before the water stress due to a reduction of *A_net_* (**Table 4**; **Fig. 3**). Over time, these returned to values matching those of uninoculated plants, suggesting a state of equilibrium between host and endophyte (**Fig. 3**). As carbon assimilates are eventually partitioned to different sinks for plant growth, tissue maintenance, or hosting microbe and endophytes, this early limitation may represent the tradeoff to establish symbiosis (Kiers & Denison, 2008; Rho et al., 2018a). At a whole-plant level, previous studies have noted this carbon expense by observing an initial suppression of growth after establishing these host-consortia symbioses in hybrid poplar (Knoth et al., 2014) and conifers (Aghai et al., 2019). Although the dynamics of carbon partitioning in plant-endophyte relationships remain largely unexplored (Rho & Kim, 2017; Kim et al., 2019), this finding supports that endophyte inoculation switches from being carbon costly to neutral.

### Inoculation alters stomatal morphology and overall patterning

We hypothesized that inoculation would change stomatal patterning and reveal a collective difference between treatments. Inoculation with the DOE mix of endophytes significantly reduced guard cell and pore length before and during the water-deficit, with less consistent effects on other individual traits (**Fig. 5**; **S3**-**S6; S12**). Conversely, across diverse plant species, reports have observed increases in stomatal length from different endophytes (Larraburu et al., 2010; Rosmana et al., 2016; Hu & Bidochka., 2021). The theory of this study assumed that certain Salicaceae endophytes may share pathways that could elicit a stomatal defense response (Carvalho et al. 2016, Melotto et al., 2017). It relied on the previous *in vivo* and *in vitro* findings that showed endophytes promote hormones involved in inhibiting stomatal development (Khan et al. 2016; Rho et al., 2018c). As our findings differ from other studies, this highlight the importance of the signals that underpin plant-microbe relationships that leads to different stomatal patterning.

Prior to the water-stress, inoculation with the DOE mix of endophyte strains showed increased stomatal numbers on both leaf surfaces (i.e., PC1; **Fig. 4**) and had a marginally significant effect on abaxial numbers post water-deficit (**Table S3**). This deviates from previous work that has shown lower abaxial stomatal densities in both inoculated rice (Rho et al., 2018c) and apple (Rho et al., 2020a). These findings may differ due to the different biological systems, experimental contexts, or methods for sample collection. For example, ABA was suggested as a potential driver of lower numbers, but several other hormones crosstalk to regulate stomatal proliferation (Wei et al., 2021). In black cottonwood, various pathways have been identified that alter stomatal density and future efforts could identify the effect of these consortia on these mechanisms (Mckown et al. 2014; Mckown et al., 2019; Chhetri 2020).

The accumulation of these changes showed an overall difference in stomatal patterning from inoculation with the DOE Mix (**Table 4**; **Table S7**). Inoculated plants tended to have denser, smaller stomata in addition to greater anatomical *g_smax_* (**Fig. 4)**. However, variation existed for each trait and overall patterning at various times along the moisture deficit (**Fig. S4 – S6; Table S8**). Inoculation explained about a fifth of the variation in overall differences between treatments (**Table S7**), suggesting its notable influence underlying changes in stomatal development. Other sources of variation between treatments reflect the impact by other environmental conditions in the greenhouse. This could be due to the different temperature or humidity experienced by the plants due to their spatial arrangement.. Further investigations should incorporate these conditions by first determining the precise effects of these on stomatal development (Bertolino et al., 2019).

While this experiment showed numerous significant effects of inoculation on host stomatal morphology (**Table 4**; Table **S3**), a follow-up greenhouse experiment showed little influence from this consortium (data not shown). We hypothesize the follow-up experiment was not successfully inoculated or that inoculation effects were overwhelmed by aspects of the environmental context such as disease pressure and low light conditions (Hendrickson 2023).

Confirming the presence of the strains used in this experiment in both control and treated plants would strengthen evidence of the relationship between endophyte abundance and plant physiology.

### Stomatal patterning corresponds to leaf gas-exchange

We hypothesized that variations in stomatal patterning would correspond to differences in *g_sw_* seen at high light. To test this, we used a regression model that represented the 13 stomatal traits, and this related to *g_sw_* under high light before (*R*^2^_*adjusted*_ = 0.24; p = 0.1) and after water-deficit (*R*^2^_*adjusted*_ = 0.63; p = 0.002). The analysis showed a significant relationship between *g_sw_* and PC1, which represented stomatal density and size (**Fig. 6**). The direction of this relationship indicated that as operational *g_sw_* increased, the magnitude of the PC scores representing plants with larger, sparser stomata also increased. The plants were distributed along a continuum, with distinct clusters based on treatment (**Fig. 6**). Specifically, uninoculated well-watered plants showed the largest, least dense stomata, resulting in the highest *g_sw_* values, while inoculated water-deficit plants displayed the lowest *g_sw_* with smaller, dense stomata.

The biomechanics of this relationship could depend on the relative importance of interconnected processes that relate structure to function. Smaller stomata can respond quicker to signals due to a greater membrane surface area to volume ratio of the guard cells (Drake et al., 2013). This would lead to a faster exchange of solutes between adjacent cells and therefore the opening and closing of the pores (Franks & Farquhar 2007). Additionally, denser stomata may indicate more tightly packed cells across the epidermis, which could increase the mechanical advantage over the guard cells, potentially limiting a greater aperture (Dow et al., 2014). This could be amplified by overlapping diffusion shells that would reduce transpiration (Lehmann & Or, 2015). However, considering only the biomechanical effect ignores physiological impacts, like osmotic shuffling or other active processes that could alter the stomatal response. The exact signals may involve the interplay between biomechanical processes, hormonal changes, and other physiological functions. Preliminary investigations of dynamic stomatal responses, following previously developed protocols (McAusland et al., 2016; Durand et al., 2019), showed relatively little variation to changing light (data not shown); therefore, future efforts may require highly controlled conditions when connecting microbial effects on stomatal responsiveness.

Building on the previous discussion, the changes in leaf physiology and morphology from inoculation may extend to whole-plant water use efficiency (WUE). Our results show that endophyte inoculation could reduce stomatal size and increase density in black cottonwood, and this related to increases in iWUE at high light at the leaf level (**Table 4**). However, instantaneous leaf-level water-use efficiency does not always translate to the whole plant scale due in part to the complex relationship between canopy architecture and microclimate (Medrano et al., 2015; Leakey et al., 2019). In previous work with Salicaceae endophytes and black cottonwood, no relationship was found between these parameters (Banan et al., 2024). This is further complicated by the uncertainty of inoculation effects associated with diurnal light levels (Rho et al., 2018c; Banan et al. 2024). One approach could integrate iWUE over the course of the day, but this still generates variable results partly because of leaf position in relation to light (Medrano et al., 2015).

Alternatively, one pathway may incorporate scaling relationships between stomata morphology and other leaf traits, such as mesophyll anatomy and specific leaf area (SLA). For example, in black cottonwood, stomatal density has shown to inversely correlate with SLA and directly with biomass accumulation (Afas et al., 2006). If this relationship holds true, inoculation may confer a unique leaf phenotype that would conserve water through less transpirational area (i.e., lower SLA) and greater biomass. Further mechanistic hypotheses could be based on these relationships, including leaf mesophyll anatomy and amphistomy (Drake et al., 2019). A morphometric approach may address these gaps and contribute to a comprehensive understanding of the inoculation effect in determining water use efficiency of the whole plant.

## CONCLUSION

This study contributes to the understanding of plant-microbe interactions, its influence on stomatal development, and its relation on leaf gas-exchange. Further, it highlights the potential for harnessing these synergistic partnerships to improve water-use efficiency in commercially and ecologically valuable trees. Forging ahead, more extensive studies that quantify endophyte strains will strengthen our understanding behind the mechanisms underlying plant-microbe interactions. These are promising advancements for enhancing water-use efficiency and sustainable productivity in forest systems, contributing to the larger efforts of alleviating the impacts of climate change.

## SUPPLEMENTARY DATA

Fig. S1. Correspondence of measured stomatal dimensions with those obtained from StomataGSmax (Gibbs et al., 2021).

Fig. S2. Violin plots of leaf gas-exchange taken light saturation (2000 μmol photons m-2 s-1) from the youngest, fully expanded leaf pre and post water-deficit.

Table S3. Effect sizes (adj η_p^2) of each treatment for the examined traits before and after water-stress.

Fig. S4. Violin plots of adaxial stomatal anatomy taken at each sampling timepoint.

Fig. S5. Violin plots of abaxial stomatal anatomy taken at each sampling timepoint.

Fig. S6. Violin plots of adaxial and abaxial stomatal densities and their ratios taken at each sampling timepoint.

Table S7. PERMANOVA tables before and after water-stress, showing the effects of inoculation, water-deficit, and their interaction.

Table S8. PERMANOVA tables from each timepoint examining the effects of inoculation, water-deficit, and their interaction on stomatal patterning.

Fig. S9. Matrices representing the correlations between each stomatal variable.

Table S10. Loading tables generated from the principal component analysis, aggregated across the entire experiment.

Table S11. Loading tables generated during the principal component analysis at each timepoint. Fig. S12. Main inoculation effect on adaxial stomatal traits from the youngest, fully expanded leaf pre and post water-deficit.

Fig. S13. Main inoculation effect on stomatal density and ratio form the youngest, fully expanded leaf pre and post water-deficit.

## ACKNOWLEDGEMENTS

Thank you to Annie Bilotta and David Zuckerman for greenhouse support, and Sabrina Zerrade for help with data processing.

## AUTHOR CONTRIBUTIONS

MH conceptualized, designed and executed the experiment, collected data, analyzed and interpreted the results, and wrote the manuscript. DB conceptualized, designed and executed the experiment, collected data, and interpreted the results. RT designed and performed the microbiological aspects of the experiment and wrote the corresponding methods. JDB helped analyze the results and edited the manuscript. SLD designed and performed the microbiological aspects of this experiment and edited the manuscript. SHK conceptualized, designed, and supervised the experiment, interpreted the results, and edited the manuscript.

## DATA AVAILABILITY

The data that support the findings of this study are openly available in Dryad at https://doi.org/10.5061/dryad.zpc866th9.

